# Topological rearrangements activate the HerA-DUF anti-phage defense system

**DOI:** 10.1101/2024.10.24.620088

**Authors:** Anthony D. Rish, Elizabeth Fosuah, Zhangfei Shen, Ila A. Marathe, Vicki H. Wysocki, Tian-Min Fu

**Affiliations:** Department of Biological Chemistry and Pharmacology, The Ohio State University, Columbus, OH 43210, USA; The Ohio State University Comprehensive Cancer Center, Columbus, OH 43210, USA; Program of OSBP, The Ohio State University, Columbus, OH 43210, USA; Center for RNA Biology, The Ohio State University, Columbus, OH 43210, USA; Department of Chemistry and Biochemistry, The Ohio State University, Columbus, OH 43210, USA; Native Mass Spectrometry Guided Structural Biology Center, The Ohio State University, Columbus, OH, 43210, USA

## Abstract

Leveraging the rich structural information provided by AlphaFold, we used integrated experimental approaches to characterize the HerA-DUF4297 (DUF) anti-phage defense system, in which DUF is of unknown function. To infer the function of DUF, we performed structure-guided genomic analysis and found that DUF homologs are universally present in bacterial immune defense systems. One notable homolog of DUF is Cap4, a universal effector with nuclease activity in CBASS, the most prevalent anti-phage system in bacteria. To test the inferred nuclease function of DUF, we performed biochemical experiments and discovered that the DUF only exhibits activity against DNA substrates when it is bound by HerA. To understand how HerA activates DUF, we determined the structures of DUF and the HerA-DUF complex. DUF forms large oligomeric assemblies with or without HerA, suggesting that oligomerization per se is not sufficient for DUF activation. Instead, DUF activation requires dramatic topological rearrangements that propagate from HerA to the entire HerA-DUF complex, leading to reorganization of DUF for effective DNA cleavage. We further validated these structural insights by structure-guided mutagenesis. Together, these findings reveal dramatic topological rearrangements throughout the HerA-DUF complex, challenge the long-standing dogma that protein oligomerization alone activates immune signaling, and may inform the activation mechanism of CBASS.

## INTRODUCTION

Utilizing the wealth of structures predicted by AlphaFold, we are witnessing a transformative shift in biomedical research towards characterizing previously enigmatic proteins^1,2^. Recent advancements in artificial intelligence-driven structural prediction have yielded numerous highly accurate three-dimensional (3D) protein models^1,3^. Given the pivotal role of 3D structures in dictating protein function, these predictions offer invaluable guidance for unraveling protein functions.

The arms race between phages and bacteria has prompted the development of diverse anti-phage defense mechanisms in bacteria^4–6^. Bioinformatic analyses have uncovered numerous bacterial immune systems, many of which harbor components with elusive functions^7,8^. One such system, HerA-DUF4297, comprises HerA, a well-characterized ATPase, and DUF4297 (DUF), a protein of unknown function^8^ (Figure 1A). Prior research has revealed that both proteins are required for effective anti-phage defense, though the mechanisms remain unknown^8^.

**Figure 1.**
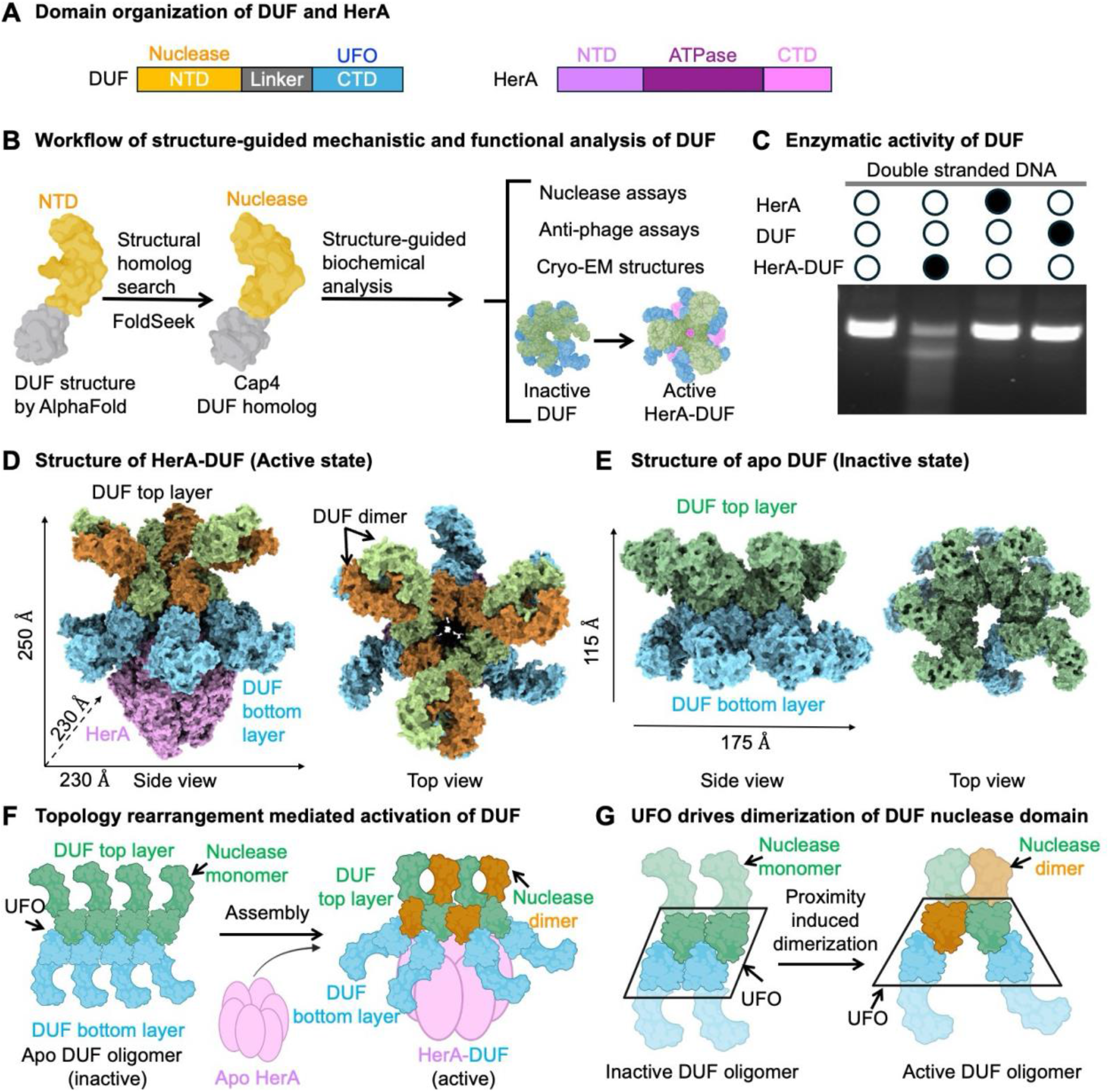
Integrated functional characterization of HerA-DUF (A) Schematic diagrams of DUF and HerA with domains color coded. (B) Workflow of integrated mechanistic and functional analysis of DUF. AlphaFold predicted structure of DUF was used to identify its structural homologs using Foldseek. Based on the functions of Cap4, a DUF structural homolog, we designed biochemical and structural experiments to elucidate functions of DUF. (C) HerA-DUF, but not DUF alone or HerA alone, displays robust nuclease activity. The gel is a representative of assays performed in triplicate. (D) Cryo-EM structure of active HerA-DUF complex from the top and side views. DUF assembles as a trimer of dimers in the top layer, while the bottom layer of DUF mediates interactions with HerA. (E) Cryo-EM structure of inactive apo DUF from the top and side views. Apo DUF assembles as an oligomer with distinct topological arrangement from active DUF. (F) UFO domain drives the formation of distinct DUF oligomers across different states. Upon forming a complex with HerA, DUF undergoes dramatic topological rearrangement, leading to the activation of DUF. (G) Upon interacting with HerA, the rearrangement of the UFO domain propagates to the nuclease domain, leading to proximity-induced dimerization of DUF nuclease domains for activation.

Here, we combine data mining, structural elucidation, and biochemical assays to characterize the function of the HerA-DUF anti-phage defense system. Leveraging the AlphaFold-predicted structure of DUF, encompassing an N-terminal domain (NTD) and a C-terminal domain (CTD), we utilized Foldseek to identify structural homologs of DUF ^9^ (Figure 1B). Remarkably, our exploration unveiled striking similarities between the NTD of DUF and the nuclease domain of Cap4, despite modest primary sequence identity^10^ (Figure 1B). Subsequent biochemical analyses revealed that DUF alone lacked nuclease activity, whereas the HerA-DUF complex exhibited robust nuclease activity (Figure 1C).

Further cryo-EM structural analysis depicts distinct topological arrangements of DUF alone and in complex with HerA (Figures 1D, 1E, and S1A-G; Table S1). While DUF alone forms compact oligomers with tightly packed nuclease domains, its direct association with HerA induces the rearrangement of DUF CTD, denoted as Universal Fold for Oligomerization (UFO) domain, leading to proximity-induced formation of three clamp-shaped dimers of DUF nuclease domains, crucial for activating DUF nuclease activity (Figures 1F and 1G).

Together, these findings highlight an integrated approach to study protein function, uncover dramatic topological rearrangements of DUF, and challenge the prior assumption that protein oligomerization by itself is sufficient to drive immune signaling.

## RESULTS

### Structure-guided clustering of DUF nuclease domain-containing proteins

To examine the structural and functional diversity of DUF homologs in bacterial immunity, we analyzed more than 1,000 representative DUF homologs, chosen for their structural homology to the nuclease domain of DUF, generated a phylogenetic tree of these proteins, and clustered them into different groups based on their sequences (Figures 2A and 2B). We used the AlphaFold predicted structure of the DUF nuclease domain as a search template to identify structural homologs with Foldseek ^9,11^. Based on the Foldseek search against AlphaFold database (AFDB), we selected 48 protein matches from AFDB, including a Cap4 endonuclease with low sequence identity of 10%, for genomic characterization of DUF homologs^10,12^. We did PSI-BLAST to identify sequence homologs of the 48 targets^13^. Eventually, we collected 1,009 DUF nuclease-containing proteins for further analysis. Phylogenetic analysis showed that these proteins can be classified into several major groups based on their primary sequences (Figure 2A). Among those major groups, a few well-characterized bacterial immune systems were identified and clustered, including the CBASS system and DEAD/DEAH box helicases^4^ (Figures 2A and 2B). In contrast, the AVAST systems are scattered in different groups, indicating their sequence divergence^8^ (Figures 2A and S2A). The HerA-DUF system was distinctly separated from these other systems, indicating a relatively distal-evolutionary relationship at the sequence level (Figures 2A and 2B).

**Figure 2.**
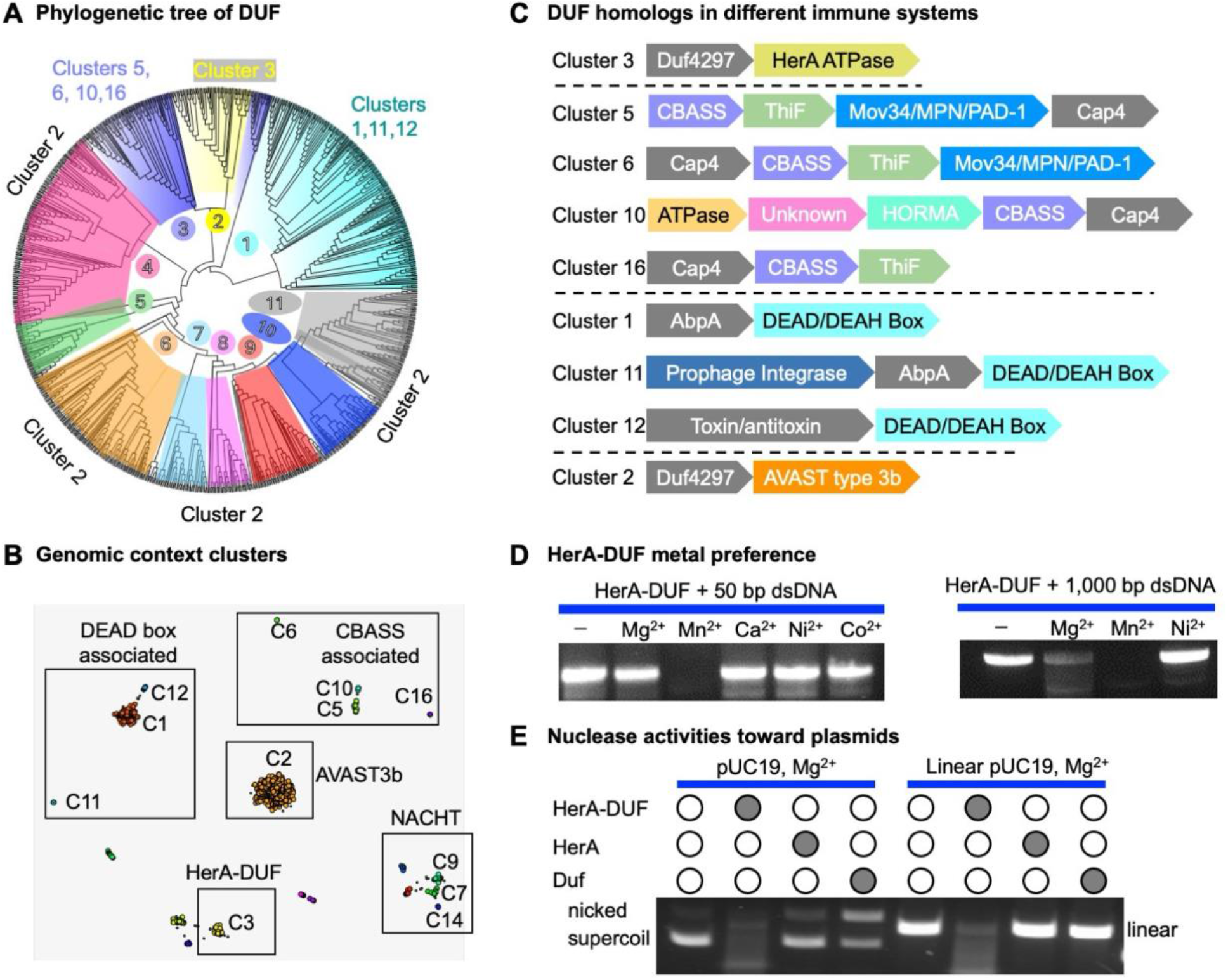
Genomic characterization of DUF by structural fold. (A) Phylogenetic tree of DUF with N = 1009 DUF nuclease domain-containing sequences with some major conserved clusters highlighted based on shared biological function. The tree is configured in a polar tree layout, rooted at the midpoint with the root hidden, nodes set in increasing order, and branches transformed in cladogram style for ease of visualization. Cluster 2 (AVAST3b family) has poorly conserved sequences and spans roughly 70% of the tree. (B) 2D genomic context cluster map, generated by GCsnap, with DUF nuclease domain-containing proteins clustered by biological function. Clusters associated with similar biological functions or protein classes are boxed together to aid visualization. (C) Cartoon representation of gene families within each identified cluster occurring with ≥ 35% frequency. Each DUF nuclease domain-containing gene is colored grey. Clusters are grouped into associated functional families by dashed lines. (D) HerA-Duf displays robust nuclease activities towards 50 bp dsDNA in the presence of Mn^2+^ but not other cations tested. In contrast, HerA-Duf can effectively process DNA substrates of 1,000 bp in the presence of Mg^2+^ or Mn^2+^ with higher cleavage efficiency for Mn^2+^. (E) HerA-DUF can eliminate plasmids in supercoiled, circular, and linear forms. In contrast, apo DUF can only nick supercoiled into circular plasmid but cannot cleave linear plasmid, indicating a low nuclease activity. The gels shown are representative of assays performed in triplicate.

Further clustering of all the DUF homologs with CLANS until equilibrium generated 18 clusters^14^ (Figure 2C). The majority of these clusters can be further grouped into five major bacterial immune systems, including CBASS, DEAD-Box helicase associated proteins, NACHT, and AVAST type 3b family of proteins (Figure 2B). Cluster 3 represents the HerA-DUF system (Figure 2C). We then explored the defense islands, or the surrounding genetic loci of these clustered sequence groups with GCsnap and we identified some conserved gene families within the clustered groups^15^ (Figure 2C). Cluster 3 did not include any other conserved families besides HerA and DUF (Figures 2C and S2A). In contrast, CBASS and DEAD Box helicase groups contained a variety of associated conserved families located within the genetic loci, underscoring diverse functional roles of DUF nuclease domain homologs in different anti-phage systems (Figure 2C).

We further analyzed the CTD of these DUF nuclease domain-containing proteins and found that they contain structurally diverse CTDs. For example, the CTD of Cap4 is a SAVED domain that can sense oligonucleotides and form an oligomer to promote the activation of the nuclease domain ^10^ (Figure 3B). In contrast, the CTD of DUF in the HerA- DUF system is structurally homologous to the large subunit of prokaryotic SPARSA SIR2 domain and the N-terminal domain of N-acetylglutamate synthase^16,17^ (Figures 3C and 3D). Like the Cap4 CTD ^10^, the CTDs of other DUF homologs potentially contribute to the activation of their nuclease domains.

**Figure 3.**
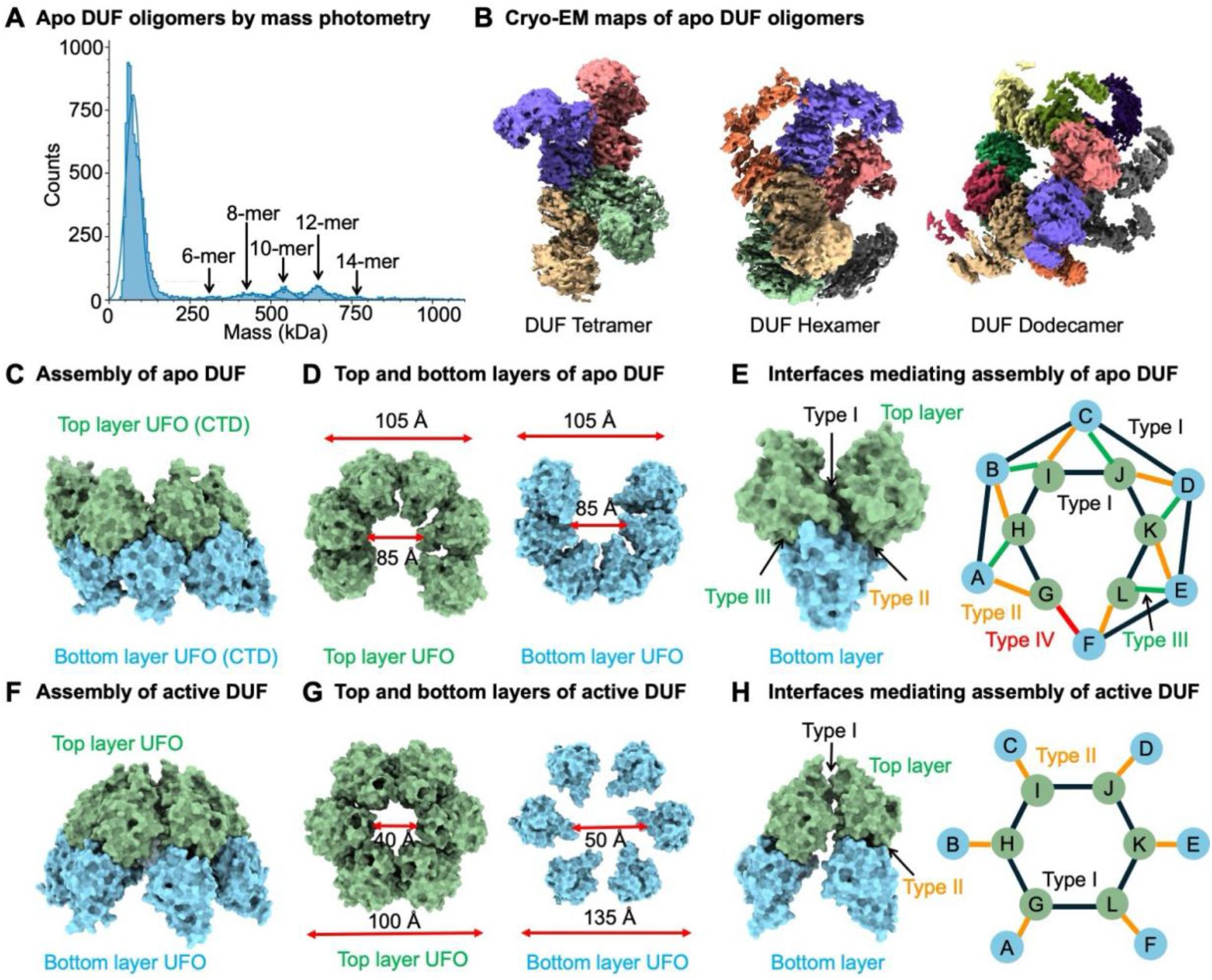
UFO drives the formation of divergent DUF oligomers. (A) DUF exists as variable oligomers revealed by mass photometry. (B) Cryo-EM maps of DUF tetramer, hexamer, and dodecamer. (C) Apo DUF assembles into a cylindric architecture with two layers, mediated by the UFO domain interactions. (D) Protomers in the top and bottom layers of apo DUF form an open-ring structure with an inner diameter of 85 Å. (E) Four types of interfaces mediate the assembly of UFO domains in apo DUF. Type I exists between adjacent subunits within each layer, while type II and type III interfaces mediate interactions between the top and bottom layers. Type IV is uniquely positioned to close the top and bottom ring. (F) Active DUF assembles into a cone-like structure with a wide bottom and a narrow top through UFO-UFO interactions. (G) The top layer of active DUF UFO domains form a closed hexameric ring with a central pore, while the bottom layer forms a dispersed hexameric ring. (H) Two types of UFO interfaces, comparable to type I and type II in apo DUF, dictate the assembly of active DUF.

Together, this analysis showed that DUF nuclease domain-containing proteins, with diverse sequences and structures, are wildly existing in diverse bacterial immune systems, underscoring their functional importance in anti-phage immune defense.

### DUF-HerA promotes nuclease activity of DUF

Given that Cap4 is a metal ion-dependent endonuclease^10^, we tested the metal preference of DUF using DNA substrates of various lengths (Figures 2D and S2E-G). We found that HerA-DUF, but not DUF alone, can effectively process short dsDNA in the presence of manganese ions, but not with other divalent cations tested (Figures 2D and 2E-G). In contrast, HerA-DUF can process long dsDNA in the presence of either magnesium or manganese. Since HerA-DUF, but not DUF alone, can process long dsDNA in the presence of magnesium (Figure 1C), we further tested whether this holds true for plasmid DNA. We found that HerA-DUF can digest plasmid DNA, whereas DUF alone only generates a nicked product, highlighting the importance of HerA-DUF complex formation in enhancing DUF nuclease activity (Figure 2E). Consistently, HerA-DUF can effectively process linearized plasmid DNA, while DUF alone fails to do so (Figure 2E). Together, these data support that HerA-DUF complex formation substantially promotes the catalytic activity of DUF.

### UFO domain drives the formation of diverse DUF oligomers

As oligomerization was required for the activation of Cap4^10^, we investigated whether DUF assembles as an oligomer. Unexpectedly, DUF alone displayed a relatively broad peak on gel filtration, indicating non-homogeneous assemblies (Figure S3A). To confirm the oligomeric status of DUF, we performed mass photometry analysis, revealing that DUF alone existed in various oligomers, including monomers, dimers, hexamers, octamers, decamers, dodecamers, and tetradecamers (Figure 3A).

Our structural analysis revealed that DUF alone indeed forms variable oligomers, consistent with our gel filtration and mass photometry findings (Figures 3B and S1B). We identified tetramers, hexamers, and dodecamers in our structural analysis with dodecamers being the dominant species (Figures S1C, S3B, and S3C). The oligomerization is mainly mediated by the CTD of DUF (Figure 3C). The CTD of DUF adopts an α-β-α sandwich structure with a six-stranded ?-sheet surrounded by six a helices (Figure S3D). As the CTD of DUF mediates protein oligomerization, we denoted this domain as a Universal Fold for Oligomerization (UFO) domain (Figure 3C).

Using the dodecamer structure of inactive DUF as an example, we found that the 12 protomers of apo DUF form a bilayer structure with six protomers in each layer, like a cylinder (Figure 3C-E). Notably, the six protomers in each layer form an open-ring structure through extensive interactions with neighboring subunits (Figure 3D). The two layers are linked together through interactions mediated by one protomer from the top layer and one protomer from the bottom layer in a shoulder-to-shoulder manner (Figure 3E and S3E). Four different interfaces are identified in the dodecamer of apo DUF, denoted as Interface I to Interface IV (Figure 3E). Interface I mediates interactions within each layer with a buried area of ∼ 500 Å² (Figures 3E and S3F). Interface II and Interface III contribute to the assembly of protomers between layers with buried areas of ∼1,260 Å² and ∼570 Å², respectively (Figures 3E, S3G, and S3H). Detailed analysis showed that all three interfaces are dominated by hydrophilic and charged residues (Figures S3F-S3H). In contrast, interface IV is mainly contributed by hydrophilic residues at the linker region with a buried area of ∼470 Å^2^, which seals the two open-rings on the top and at the bottom layers (Figures S3E and S3I).

In its active state, DUF also assembles as a dodecamer through UFO-UFO interactions (Figures 3F-3H). Similar to the apo structure, the 12 protomers of DUF in the active state form a bilayer structure with six protomers in each layer (Figures 3F and 3G). However, unlike the open-ring structure of the apo state, the six protomers in the top layer form a closed ring through extensive interactions with neighboring subunits, while the six protomers in the bottom layer do not interact with each other (Figures 3G-3H). Consequently, the UFO domains in active DUF resemble a cone, starkly contrasting the cylindrical structure observed in the inactive state (Figures 3C and 3F). These topological differences arise from changes in the interacting interfaces. In active DUF oligomers, two distinct interfaces, corresponding to Interfaces I and II in the apo state, are observed (Figures 3H, S3J, and S3K). Interface I, with a buried area of ∼470 Å^2^, mediates interactions between neighboring subunits in the top layer, while Interface II, with a buried area of ∼830 Å^2^, facilitates interactions between the top and bottom layers (Figures 3H, S3J, and S3K). Furthermore, the total buried areas in the active DUF are substantially smaller than those in the apo state. These differences dictate the topological variations of UFO in active versus apo DUF, which propagate to the nuclease domain, leading to the topological divergence of the nuclease domains between active and apo DUF.

Assembly of the UFO domain is critical for the activation of DUF. Substitution of interfacial residues can effectively disrupt the oligomerization of DUF as determined by gel filtration analysis (Figure S3L). These interfacial mutants substantially reduced the nuclease activity of the HerA-DUF complex and their activities for anti-phage defense (Figures S3M and S3N).

Together, our structural analysis underscores the critical role of the UFO domain in driving the divergence of DUF oligomers across different states.

### Topological rearrangements activate DUF

The topological arrangement of active DUF exhibits dramatic differences from that of apo DUF, revealing the process of DUF activation (Figures 4A and 4B). In the apo DUF dodecamer, protomers in the top and bottom layers are orientated clockwise and counterclockwise to each other, leading to the tight packing of the nuclease domains of DUF (Figure 4A). In contrast, the conformations of the DUF protomers in active state undergo a global rearrangement, revealing a mechanism for the activation of DUF. First, the bottom layer of active DUF displayed a dispersed conformation (Figure 4B). Second, the neighboring protomers of DUF on the top layer form a swapped dimer (Figure 4B). In total, there are three pairs of DUF dimers on the top layer. These dimers can form between any adjacent DUF protomers on the top layer, resulting in two possible configurations of three pairs of dimers (Figure S1G). Together, these data revealed a mechanism of global domain rearrangement during the switching of DUF from apo state to active state.

**Figure 4.**
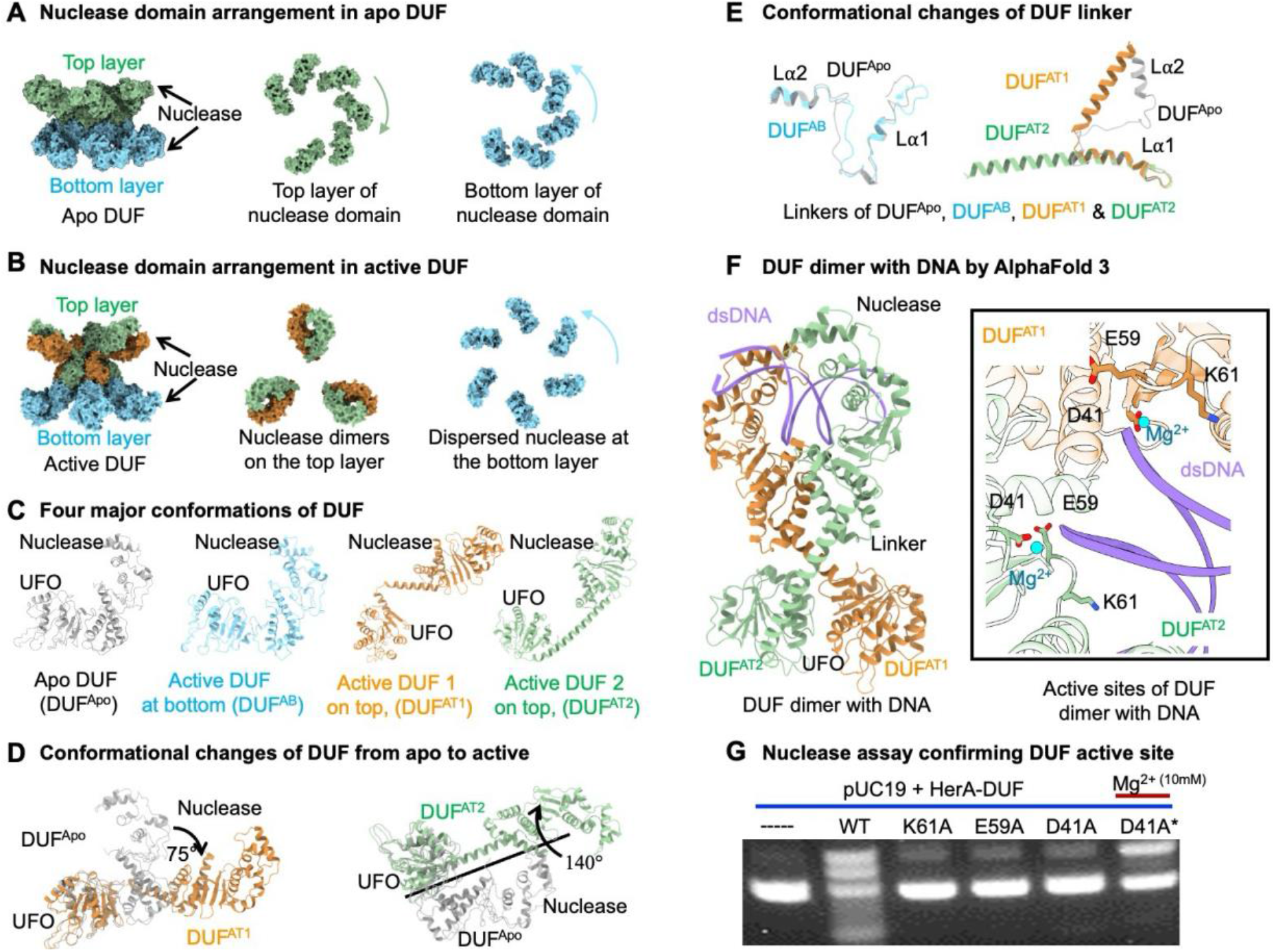
Topological rearrangement of nuclease domain activates DUF. (A) DUF nuclease domains in each layer of apo DUF rotate anti-parallel to each other and pack asymmetrically without contacting each other. (B) Nuclease domains in each layer of active DUF display distinct rearrangements. In the top layer, two adjacent nuclease domains (orange and green) form a dimer. In contrast, nuclease domains in the bottom layer (blue) adopts a dispersed ring-like structure. (C) All the DUF protomers in apo state share the same conformation, while there are three conformations observed in active DUF protomers. (D) Overlaid protomers of DUF in apo and active state, revealing their domain rearrangement across states. (E) Conformation changes in the linker region of DUF drive the nuclease domain rearrangement of DUF. (F) Structure of DUF dimer in complex with DNA (purple) and a blowup of the DUF active site with magnesium, predicted by AlphaFold 3. (G) Substitution of catalytic residues abolished nuclease activity of HerA-DUF. D41A* represents a rescuing experiment by adding additional magnesium, indicating that D41 is responsible for coordinating magnesium. The gels shown are representative of assays performed in triplicate.

The topological rearrangement of DUF is driven by rigid body rotation of the nuclease domain, accompanied by pronounced structural changes in the linker region between the nuclease domain and the UFO domain (Figures 4C-4E and S4A-C). The protomer of apo DUF adopts a folded conformation (denoted as DUF^Apo^), whereas protomers of active DUF have three conformations (Figure 4C). As such, we denoted protomers at the bottom layer as DUF^AB^, protomers on the top layer as DUF^AT1^ and DUF^AT2^ due to their conformational differences in the linker region. The conformation of DUF^AB^ resembles the conformation of DUF^Apo^ with limited tilting in the nuclease domain (Figure S4A). However, when aligned with DUF^Apo^ using the UFO domain, the nuclease domains of DUF^AT1^ and DUF^AT2^ rotate about 75° and 140°, respectively, along an axis at the junction between the nuclease domain and the UFO domain for two neighboring protomers on the top layer (Figure 4D). The structural change is approximately a rigid body rotation as the UFO domain and the nuclease domain align well in gross conformations between the inactive and the active states (Figures S4B and S4C). Strikingly, the linker region between the nuclease domain and the UFO domain in protomers of the top layer of active DUF undergoes dramatic conformational changes, compared to that in the apo DUF (Figure 4E). The conformational changes in the linker region include secondary structure changes from loops into alpha helices and changes from folded conformation to a stretched conformation (Figure 4E). For the protomers at the bottom layer of active DUF, the linker region adopted a similar conformation with that in the inactive DUF but titled a certain angle, leading to an extended conformation of the nuclease domain (Figures 4E and S4A). Together, these data revealed the structural basis of DUF topological rearrangement from an inactive state to an active state.

Dimerization of the DUF nuclease domain is likely crucial for binding and cleaving DNA substrates. This dimerization is mediated not only by the UFO domains but also by linker-linker and nuclease-nuclease interactions, with a topology unlike that of the nuclease domain dimer of AVAST3 ^18^ (Figures 4F, S4D and S4E). Hydrophilic residues dominate the interactions in the linker region, while hydrophobic residues facilitate the interactions between neighboring nuclease domains (Figures S4F and S4G). Together, these interactions create a buried interface of ∼1,500 Å², stabilizing the dimeric DUF nuclease domain for effective DNA binding. The AlphaFold3-predicted structure of the DUF-DNA complex reveals that double-stranded DNA is coordinated in the center of the clamp-shaped DUF dimer^2^ (Figure 4F). Key residues, including hydrophilic and positively charged ones, are critical for this coordination. Consequently, the dsDNA is well-coordinated by the clamp-shaped nuclease domain, with two strands poised for processing by the two protomers, explaining the robust activity of the DUF-HerA complexes (Figure 4F).

The active site of the DUF nuclease domain is composed of a catalytic triad, resembling that of the Cap4 nuclease domain ^10^ (Figure 4F). Based on the AlphaFold3 prediction^2^, D41 is responsible for coordinating magnesium while catalysis also requires E59 and K61 (Figure 4F). Substitutions of these residues abolished the nuclease activity of DUF, underscoring the functional importance of these residues in catalysis (Figure 4G).

Together, these results uncover the mechanism of DUF activation driven by the global conformational changes of the DUF nuclease domain.

### Interactions between HerA and DUF activate DUF

Interactions between HerA and DUF are mediated by the HAS domain of HerA and the UFO domain of DUF (Figures 5A-D). HerA is composed of three domains: HAS, a RecA ATPase domain, and an inserted helical bundle domain, followed by a C-terminal brace (Figures S5A and S5B). Notably, the HerA we studied contains a long C-terminal brace, contrasting with the HerA molecule in the HerA-Sir2 complex ^19,20^ (Figure S5C). The cryo-EM structure of the HerA-DUF complex reveals that 12 copies of DUF assemble into two layers on top of the HerA hexamer (Figure 5A). The bottom layer of DUF, but not the top layer, forms extensive interactions with the HerA hexamer, with a total buried interface of

**Figure 5.**
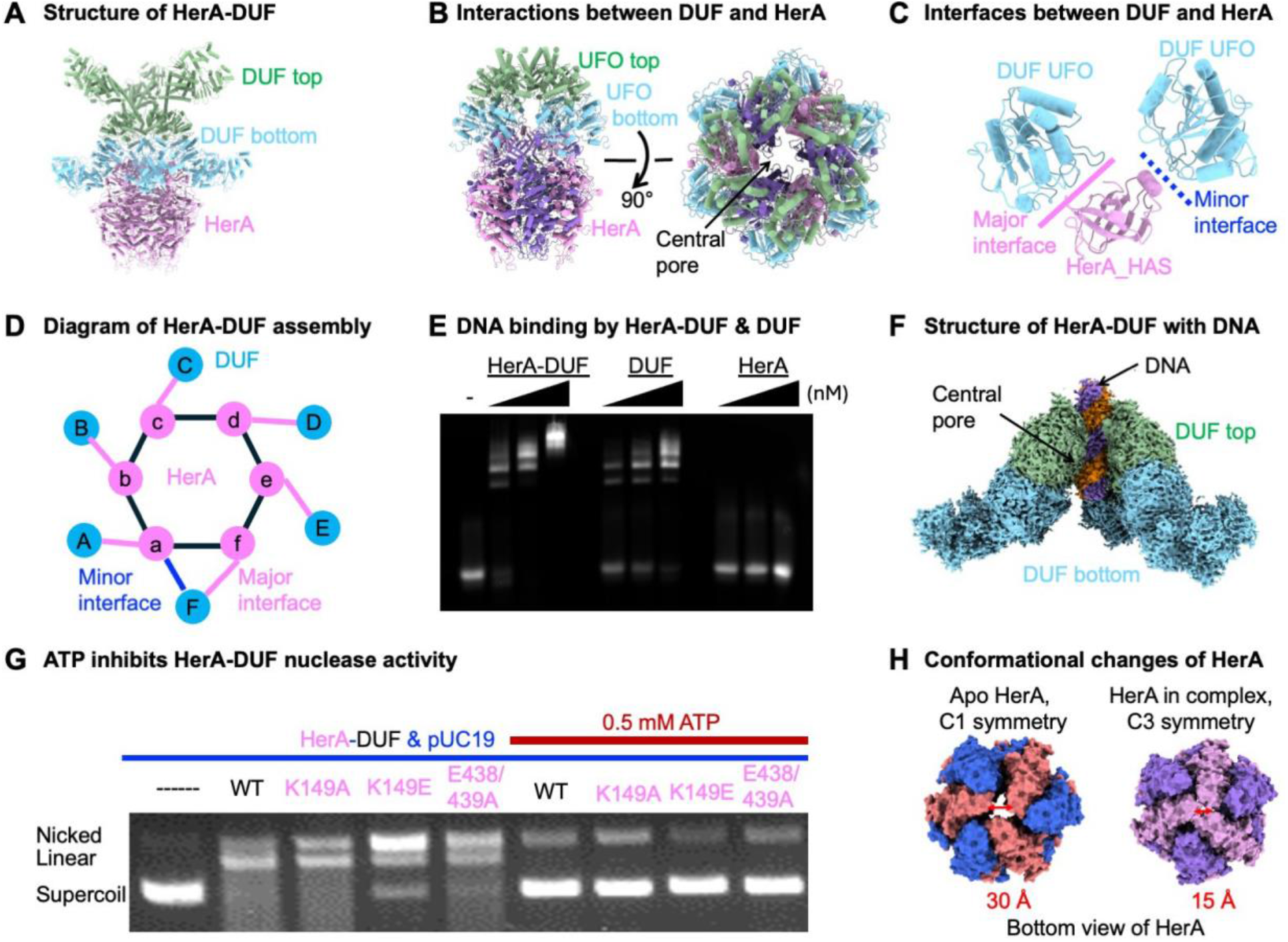
Interactions between HerA and DUF activate DUF. (A) Structure of HerA-DUF complex in ribbon diagram with HerA and DUF color coded. (B) Top view and side view of HerA in complex with DUF UFO domains. A central channel runs through the entire HerA-DUF complex with a large opening at the top of DUF. (C) Interfaces mediating interactions between HerA HAS domain (magenta) and DUF UFO domain (cyan). All six protomers of HerA interact with DUF through a major interface. However, only one protomer of HerA out of six interacts with a secondary UFO of DUF through a minor interface. (D) Diagram illustrating interactions between HerA and DUF. The six protomers for HerA are denoted ”a-f” and the six protomers of bottom layer DUF are denoted “A-F.” (E) The HerA-DUF complex likely binds dsDNA more efficiently than apo DUF, as revealed by gel shift assay. In contrast, HerA alone appears not to bind dsDNA. The gel shown is representative of assays performed in triplicate. (F) Cryo-EM density of DUF-DNA complex, revealing that DNA binds to the central pore of DUF. (G) ATP appears to inhibit HerA-DUF nuclease activity independent of HerA ATPase activity. The gel shown is representative of assays performed in triplicate. (H) Upon forming a complex with DUF, HerA undergoes dramatic rearrangement in comparison to the apo HerA.

∼3,100 Å² (Figures 5B and 5C). Specifically, HerA uses its N-terminal HAS domain to interact with the UFO domains of the bottom DUF layer, similar to other HerA proteins that engage partner proteins via their HAS domains (Figures 5B-5D). However, compared to the HAS domain in other HerA proteins, we found unique inserted motifs and different interaction hotspots in our case (Figures S5D and S5E). In the DUF-HerA complex, two interfaces—major and minor—mediate interactions between the HerA HAS domain and the DUF UFO domain (Figures 5C and 5D). The major interface facilitates paired interactions between all the protomers at the bottom layer of DUF and the six protomers of HerA (Figure 5C), while the minor interface occurs between one HerA protomer out of six and one DUF protomer out of six (Figure 5D). The major interface, with a buried surface area of ∼480 Å^2^, is formed by loops from both the HAS domain and the UFO domain, whereas the minor interface of ∼240 Å^2^ involves a short a-helix of the HAS domain and loops of the UFO domain (Figures 5C, S5F, and S5G). Both interfaces are dominated by hydrophilic and charged residues (Figures S5F and S5G).

Interactions between HerA and DUF induce global rearrangement of DUF, facilitating its nuclease domain dimerization and DNA binding, thus activating DUF. The unique interaction mode between HerA and DUF positions HerA between two neighboring DUF protomers in the bottom layer, preventing their interaction and creating a wider platform for the top layer (Figures 5B-5D). The top layer interacts with the bottom layer through the Type II interface and forms strong intra-layer interactions (Figure 3H). This substantial rearrangement brings the linker regions of top-layer protomers into proximity by angling the top layer UFO domains towards the central pore, facilitating the dimerization of DUF’s nuclease domains (Figures 1G and 4B). Additionally, this rearrangement aligns positively charged residues in the central pore of active DUF, facilitating DNA binding, as supported by DNA binding assays (Figures 5E, S5H, and S5I). Cryo-EM structures of the DUF-HerA complex with dsDNA reveal that the DUF UFO domain coordinates dsDNA (Figures 5F and S10 J-M; Table S2), suggesting that the UFO domain captures dsDNA and hands it to the nuclease domain for cleavage.

ATP functions to inhibit the nuclease activity of the HerA-DUF system. Our biochemical analysis reveals that 0.5 mM ATP is sufficient to inhibit the nuclease activity of the HerA-DUF system (Figure 5G). ATP hydrolysis by HerA appears to be decoupled with the nuclease activity of DUF in the system. In particular, the HerA-DUF complex formed by wild-type HerA and ATP hydrolysis defective HerA displayed similar nuclease activity and can be inhibited by 0.5 mM ATP similarly (Figure 5G and S5N). As such, there is a possibility that ATP may inhibit the activity of DUF through directly acting on DUF.

Interactions between DUF and HerA also reshaped the topological rearrangement of HerA (Figures 5H and S6A-F). To uncover conformational changes between apo HerA and HerA in the complex, we further determined the cryo-EM structure of HerA alone, revealing that HerA alone also forms a hexamer (Figures S6A-C; Table S2). Individual promoters from apo HerA and the HerA in the complex resemble each other (Figure S6G). However, they assemble into hexamers with distinct conformations. Structural comparison between apo HerA and HerA in the complex revealed four distinguished features.

First, the HerA in the complex has a C3 symmetry while the apo HerA is asymmetric (Figure 5H).

Second, the CTD of HerA in the complex packed tightly to each other with a narrow pore with a diameter of ∼15 Å in the middle (Figure 5H). In contrast, the CTD of apo HerA are distal to each other with a wider pore of ∼30 Å in diameter (Figure 5H).

Third, the Has domain in the complex form a wider pore with a diameter of 40 Å, compared to a narrower pore with a diameter of 20 Å in the apo HerA (Figure S6E).

Fourth, HerA protomers in the HerA-DUF complex form more extensive interactions with each other with a total buried surface area of ∼19,940 Å^2^, compared with a total buried area of 18,120 Å in the apo HerA hexamer (Figure S6F).

The assembly of HerA-DUF modulates the ATPase activity of HerA (Figure S6G). HerA alone displayed a modest ATP hydrolysis activity. When forming a complex with DUF, the ATPase activity of HerA was substantially promoted (Figure S6H). Perhaps, the topological rearrangement of HerA, when forming a complex with DUF, contributed to the ATP hydrolysis.

Together, the assembly of HerA-DUF leads to the global rearrangements of both DUF and HerA, promoting the nuclease activity of DUF and ATPase activity of HerA.

## DISCUSSION

In this work, we introduced a new paradigm for studying the HerA-DUF anti-phage defense system using integrated approaches, emphasizing the intricate interplay between protein structure, assembly, and activity regulation (Figure 1). Empowered by artificial intelligence, we leveraged the AlphaFold-predicted structure of DUF as a template to identify structural homologs of DUF and discovered that DUF shares a similar nuclease domain with Cap4^10^. Our genomic analysis revealed that many DUF nuclease domain-containing proteins, which contain a common nuclease domain along with a variable additional domain, are prevalent in bacterial immune systems, underscoring their importance in anti-phage defense (Figure 2). Based on this structural similarity, we hypothesized that DUF functions as an endonuclease and found that DUF displayed robust nuclease activity against various DNA substrates only in complex with HerA. To understand how HerA unleashes the nuclease activity of DUF mechanistically, we determined cryo-EM structures of apo DUF and the HerA-DUF complex, revealing that dramatic topological rearrangement of DUF oligomers drives the activation of DUF Figures 1, 3, and 4).

Our structural analysis reveals that topological rearrangements propagate across domains in the HerA-DUF complex, providing remarkable mechanistic insights. In the apo DUF state, 12 protomers assemble into a bilayer cylindrical architecture through UFO- UFO interactions, with all nuclease domains packed together as a monomer (Figure 3). In contrast, in the HerA-DUF complex, two layers of DUF stack atop a HerA hexamer, forming a cone-like structure (Figure 4). The interaction between HerA and DUF is mediated by the HAS domain of HerA and the UFO domain of the bottom DUF layer. The HAS domain positions itself between two neighboring UFO domains of DUF, disrupting the interactions between neighboring DUF protomers in the bottom layer and resulting in a dispersed configuration. This rearrangement in the bottom DUF layer propagates to the top layer, causing the UFO domains in the top layer to angle toward each other (Figure 4). This proximity induces dimerization of the nuclease domains with neighboring partners in the top layer (Figure 4). This dimerization enhances the binding and cleaving efficiency of DNA substrates, explaining the substantially higher activity of the HerA-DUF complex compared to DUF alone. Furthermore, in comparison to inactive DUF oligomers, active DUF oligomers undergo dramatic rearrangements primarily through rigid body rotations, without significant conformational changes in their UFO and nuclease domains (Figure 4).

Our study demonstrated that higher-order oligomerization of proteins is necessary but not sufficient for activating the HerA-DUF system. Previous research suggested that higher-order assemblies alone drive immune signaling^21,22^, a long-standing dogma in the field. Our study challenges this concept, revealing that protein oligomerization alone does not activate DUF. In fact, both DUF alone and the HerA-DUF complex assemble as large oligomers with distinct topological arrangements. It turns out that the topological arrangement of these assemblies dictates the activation of the HerA-DUF system. These insights could broadly extend to other systems where oligomerization by itself is insufficient for activation.

## Methods

### Genomic analysis of DUF4297

The amino acid sequence of DUF4297 from *Bacillus ap.* HMH5848 was used for protein BLAST (blastp) against standard and experimental databases^23^. 1,291 sequences were identified with *E* value 0.05 – 10^-146^ (0.0), but no CAP4 protein homologs were among the hits. A structural search with Foldseek (3Di/AA) using the atomic coordinates of *Bs*DUF NTD revealed AbpA (UniProt P52127) and Cap4 (UniProt C0VHC9) as the only probable matches with lengths close to the length of the NTD in the AFDB-Proteome or AFDB-SWISSPROT databases ^9,12^. Therefore, 48 sequences with probability > 0.7 and similar sequence lengths to the query were randomly selected from the AFDB50 database and were aligned with MUSCLE ^24^. The resulting alignment was used for PSI-BLAST (nr_pro_Dec14) with eight rounds through the MPI Bioinformatics toolkit as of Jan 2024 (BLOSUM45) ^13,25,26^. All collected sequences were filtered to a maximum sequence identity of 95% and 95% sequence coverage with MMseqs2 and clustered on the basis of blastp all-against-all pairwise searches with CLANS until equilibrium at *E* value of 1 × 10^−80^ ^14,27^.

A phylogenetic tree was created from 1009 sequences using iqtree2 (v2.2.2.6-MacOSX), with standard model selection followed by tree inference. 1500 Ultrafast bootstrap analysis with 1500 alerts and nstop 500 were used as cutoffs for tree generation. The tree is configured in a polar tree layout, rooted at the midpoint with the root hidden, nodes set in increasing order, and branches transformed in cladogram style for ease of visualization.

The resulting sequence similarity network was processed with machine learning as previously described by Durairaj, Janani, et. al. 2023 ^28^. In short, the similarity network was used as input for GCsnap (v.1.0.10.9) for the analysis of the conservation of the genomic contexts encoding for each of the proteins in the individual clusters ^15^. A window of four flanking genes was used. MMSeqs2 was used for protein family clustering ^27^, and clusters of similar genomic contexts were detected using the operon_cluster_advanced method, which uses PaCMAP (v.0.7.0)^29^ to project genomic contexts in two dimensions on the basis of their family composition and DBSCAN^30^ (as implemented in scikit-learn v.1.4.0^31^) to identify clusters of similar genomic contexts ^32–34^. Only families that were found in at least 35% of all genomic contexts were considered to avoid false positive hits. Structural models of DUF4297 homologs were built using AlphaFold (v2.3.2)^35^ and were visualized in ChimeraX (v1.7.1)^36^.

### Protein expression and purification

HerA and DUF4297 from *Bacillus ap.* HMH5848 were synthesized into a pET-Duet-1 expression vector with a C-terminal 6x histidine tag, and a pET-28a(+) expression vector with an N-terminal 6x histidine tag, respectively by GenScript, with codon optimization. Mutagenesis of HerA: K149A, K149E, and E438/439A and DUF: D41A, E59A, K61A, R337E, and D412R mutants were created with Q5 site-directed mutagenesis kit (NEB, catalog # E0554S), and verified with Sanger sequencing. See Supplementary Table 2 for primer and plasmid sequences. HerA and DUF were recombinantly expressed using *E. coli* BL21(DE3) RipL strain. Bacterial cells were grown at 37 °C in LB supplemented with 100 µg mL^−1^ ampicillin or 50 µg mL^−1^ kanamycin until they reached an OD_600_ of 0.8. HerA- DUF complex was co-expressed in LB containing ampicillin and kanamycin. Expression of protein was induced by the addition of 0.3 mM isopropyl β-D-1-thiogalactopyranoside (IPTG), and cultures were further incubated at 18 °C for 16 h. Cells were pelleted at 2000 × *g* for 20 min at 4 °C, resuspended in 20 mM Tris-HCl pH 7.5, 750 mM NaCl lysis buffer, disrupted by sonication, and the resulting lysate was centrifuged at 40,000 × *g* for 60 min at 4 °C. The supernatant was loaded onto a column containing Ni-NTA beads (Qiagen, catalog no. 30210) that were pre-equilibrated with lysis buffer. The column was washed with 10 column volumes (CV) of 20 mM Tris-HCl pH 7.5, 750 mM NaCl, 30 mM imidazole wash buffer, followed by elution using 5 CV of 20 mM Tris-HCl pH 7.5, 750 mM NaCl, 300 mM imidazole elution buffer, in 1 mL aliquots. Eluted protein was pooled and concentrated to 1 mL using an Amicon ®Ultra 50,000 NMWL centrifugal filter (Millipore, SKU UFC903024), before application to a Superdex 200 10/300 GL Increased or Superose 6 Increase 10/300 GL column (Cytvia, GE28-9909-44, GE29-0915-96) to purify protein to homogeneity. Protein was eluted with gel filtration buffer (20 mM Tris-HCl pH 7.5, 150 mM NaCl) on an ÄKTA go FPLC (Cytvia). HerA and DUF were verified by SDS-Page gel, stained with Coomassie blue. HerA-DUF (D41A) was incubated with blunt dsDNA for 30 minutes before freezing on grids.

### Cryo-EM data collection

Quantifiol R 1.2/1.3 Au 400 mesh 2 nm Carbon (Quantifoil) were glow discharged at 0.2 atm for 30 s and were loaded onto a FEI vitrobot (Thermo Fisher). 3 µL of the sample was applied to the grid face before blotting for 4 s and plunge freezing in liquid ethane. Grids were stored in liquid nitrogen prior to screening on a 200 kV Glacios (Thermo Fisher Scientific) equipped with a K3 bioquantum detector (Gatan, Inc.). Grids were again stored in liquid nitrogen until data could be collected using a 300 kV Titan Krios (Thermo Fisher Scientific), also equipped with a K3 detector. For HerA-DUF and apo DUF protein structures, 6879 movies were collected, and for apo HerA, 4,851 movies were collected using data acquisition software EPU (v2.12.1.2782REL), and TIA (v5.0 SP4) (Thermo Fisher Scientific), and 6,216 HerA-DUF (D41A)-dsDNA movies were collected using SerialEM data collection software, all at a magnification of ×81,000, a pixel size of 1.07 Å, at a total dose of 50 e^−^/Å^2^, and defocus range -0.5 to -2.0 µm.

### Cryo-EM data processing

All data processing, including patch motion correction and patch contrast transfer function (CTF) estimation, was performed in cryoSPARC (ver. 4.4.1) ^37^. The details of data processing are illustrated in supplemental data (Figures S1, S2, S5, and S6).

### Model building and refinement

The initial models of HerA and DUF4297 were predicted with AlphaFold^35^, and fitted into cryo-EM maps using ChimeraX^36^. Manual adjustments were made with refinement in Coot (0.9.8.7)^38^ and real-space refinement was performed in Phenix (v1.21-5207)^39^. Top layer DUF NTD density was resolved from noise using DeepEmhancer (v0.14)^40^ in post-processing. Structural models were validated with MolProbity (v4.5.2)^41^. All structural images were generated using ChimeraX.

### ATPase assay

The ATPase activity was measured using the ATPase/GTPase Assay Kit (Sigma-Aldrich, #113CB04A30). A working solution of 4 mM solution of ATP was prepared. Varying concentrations of the proteins (50 to 250 nM) were incubated with 1mM of ATP in the buffer of 20 mM Tris pH 8.0, 75 mM KCl, and 2 mM MgCl_2_ (included in the kit) in a 40 ml reaction at 37 °C for 30 mins. The reaction was stopped, and the Biotek Synergy HT microplate reader was used to read the reaction products at 620 nm. A phosphate standard curve was generated following the protocol from the kit. The generated standard curve was used to determine ATPase activity.

### Nuclease assays

Nuclease assays were adapted from previous work ^10^. In detail, the DNA digestion or cleavage activity of the proteins was performed by incubating 50 nM protein with 10 ng ml ^-1^ pUC19 plasmid (NEB N3041S), in a 20 µl reaction at 37 °C for varying time conditions in 20 mM Hepes pH 8.0, 75 mM NaCl, 2 mM MgCl_2_, and or ATP buffer. The reactions were stopped by adding 6x purple loading buffer (B7024S NEB). 20 µl of the samples were run and separated on a 0.5% TAE agarose gel, stained with ethidium bromide. Gels were run at 100 V for 20 min and visualized on Sapphire biomolecular imager (Azure Biosystems).

With the short DNA substrate, 400 nM of proteins were incubated in reaction buffer (20 mM Hepes pH 8.0, 75 mM NaCl, and 2 mM MgCl_2_) with 800 nM Cys-labeled nucleic acids substrates (20 or 50 overhang dsDNA, and 20 or 50 blunt dsDNA,) at 37 °C for 30 mins. The reactions were stopped with EDTA and SDS and samples were separated on 12% DNA PAGE or 2.5% TB agarose gel in TBE or TB buffer at 100 V for 30 min and visualized on Sapphire biomolecular imager (Azure Biosystems). See Supplementary Table 3 for DNA substrate sequences used.

### Helicase assays

For the helicase assay, varying concentrations of HerA (50 to 200 nM) were incubated with short DNA substrates mentioned earlier in 20 mM Hepes pH 8.0, 75 mM NaCl, 2 mM ATP, and 2 µM MgCl_2_ buffer at 37 °C for 30 mins. The reaction was quenched with EDTA and SDS and separated on 2.5% TB agarose gel.

### DNA Binding Assay (EMSA)

For fluorescent protein-DNA binding assay, protein concentrations of HerA-DUF complex, DUF, and HerA between 100 to 800 nM were incubated with 800 nM Cy5 -labeled dsDNA substrate in reaction buffer (25 mM Hepes, pH 8.0, 75 mM NaCl, and 5 mM MgCl2) at 37 °C for 15 minutes. The samples were separated on 2.5% agarose gel in 0.5% TB buffer. The gel was imagined on by Sapphire biomolecular imager (Azure Biosystems).

### Plaque assay

Plasmids containing HerA, DUF, or HerA-DUF complex or mutants were transformed into BL21(DE3) RipL cells as described in protein expression and purification. A single colony was grown in 2 mL LB containing the appropriate antibiotic, shaking at 220 rpm at 37C until OD_600_ of 0.1 before inducing with 0.2 mM IPTG and allowing to grow until OD_600_ of 0.4. Then, 500 µl of cultured bacteria was mixed with 14.5 mL 0.5% top agar at 40°C and poured onto plates containing antibiotic and 0.1 mM IPTG. After allowing the top agar to solidify, plates were serially spotted with 2.5 µl T4 phage from 10^0^ – 10^-8^. Plates were left to incubate overnight at 30°C.

### Mass photometry

To determine the stoichiometry of BsDUF, BsHerA and BsDUF + BsHerA, mass photometry (MP) experiments^42^ were performed using Refeyn TwoMP mass photometer. Glass coverslips (Thorlabs, CG15KH) needed for the measurements were cleaned by rinsing several times alternatingly with water and 100% isopropanol and drying with nitrogen. Culture well-containing silicone gaskets (Sigma, GBL103250-10EA) were cleaned similarly and attached to the glass coverlips. A droplet of index-matched immersion oil (Zeiss Immersol 518 F, UFI: 4Y00-R0DY-1007-3VF3) was added on the mass photometer objective before placing the coverslip on top. All measurements (calibrants and analytes) were carried out in 20 mM HEPES (pH 7.4), 150 mM NaCl, 0.5 mM TCEP. Data was acquired using AcquireMP^TM^ and analyzed using DiscoverMP^TM^. Each mass photometer movie was recorded for one minute. For calibration, an equimolar mixture of β-amylase (Sigma: A87811VL) (monomer = 56 kDa, dimer = 112 kDa and tetramer = 224 kDa) and thyroglobulin (Sigma, T9145) (monomer = 330 kDa and dimer = 660 kDa) was used. Calibration was obtained by plotting ratiometric contrast of each gaussian distribution against the known masses of calibrants and had an R^2^ > 0.999. Prior to MP measurements, all analytes were diluted to 200 nM. This diluted stock was then used for measurements. Final concentration of samples in gasket wells was 8-12 nM. Masses of the following samples were measured: 1) BsDUF alone, 2) BsHerA alone, 3) BsDUF + BsHerA. Each measurement was carried out in triplicates. Masses were determined using the calibration obtained with β-amylase and thyroglobulin.

### Statistical methods

Excel and GraphPad PRISM were used for statistical analysis. Standard t-tests were used to quantify differences of ATPases activity between HerA and HerA-DUF. Error bars are defined in figure legends.

## Acknowledgments

We thank Dr. Shiyu Xia at California Institute of Technology, Drs. Chen Wang and Wen Tang, Jiale Xie, Benjamin Pastore for discussion and critical comments for the manuscript. Grids screening was performed at OSU CEMAS with the assistance of Drs. Giovanna Grandinetti and Yoshie Narui. Cryo-EM data were collected at NCI cryo-EM national centers supported by grants from the NIH National Institute of Health Common Fund Transformative High Resolution Cryo-Electron Microscopy program. A.D.R was supported by an NIH T32 grant (GM 118291-05 and GM144293-01), and the OSU Presidential Fellowship, and T.M.F by a grant from the NIH National Institute of General Medical Sciences (GM103310). I.A.M and V.H.M. are supported by NIH HHS 1RM1GM149374.

## Author contributions

T.M.F. conceived the project. A.D.R. and E.F. performed molecular cloning and biochemical purification. A.D.R. performed genomic analysis, determined the cryo-EM structures, and built the models. E.F., and A.D.R. made all the mutants, did the biochemical assays and phage plaque assay. I.M. did all the mass photometry experiments with the supervision of V.H.W. T.M.F., A.D.R., E.F., and Z.S. analyzed the data. T.M.F. and A.D. R. prepared figures and wrote the manuscript with inputs from all the authors.

## Competing interests

All authors declare they have no competing interests.

## Data availability

Accession numbers for DUF in apo state, the DUF-HerA complex, and HerA alone are as follows: (coordinates of atomic models:9C1M, 9C1N, 9C5X, 9C1O, and 9C1X, deposited to Protein Data Bank), and (density map: EMD-45124, EMD-45234, EMD-45126, EMD- 45132 deposited to Electron Microscopy Data Bank). Density maps of DUF tetramer, hexamer, and dodecamer, HerA-DUF in complex with DNA were deposited to Electron Microscopy Data Bank with codes of EMD-45129, EMD-34133, EMD-45134, and EMD- 45280, respectively. All data needed to evaluate the conclusions in the paper are present in the paper.

## Notes

### Competing Interest Statement

The authors have declared no competing interest.

